# Visual cues and strategies for perceiving elasticity

**DOI:** 10.1101/2023.03.24.534031

**Authors:** Vivian C. Paulun, Florian S. Bayer, Joshua B. Tenenbaum, Roland W. Fleming

## Abstract

A key to interacting with the physical world is the ability to visually infer object properties, like elasticity, allowing us to anticipate object behavior. Such perceptual inferences continue to challenge AI systems, highlighting the complexity of the underlying computations. How does the human brain solve this task? Here, we propose a resource-rational model based on learned statistics of object motion to explain how humans judge elasticity. We created 100,000 physics-based simulations of bouncing cubes with different elasticities and found that even tiny changes in initial conditions (e.g., orientation) yield starkly different trajectories. Yet, across these simulations, we identified 23 motion features that capture natural variations in elasticity. Although a weighted combination of these features reliably predicts physical elasticity, surprisingly, humans do not seem to employ cue combination when judging elasticity. Instead, they switch between different cues. A series of experiments designed to carefully tease apart several competing heuristics, suggests that observers switch between different computationally efficient yet informative cues depending on the information available in the stimulus.

## Introduction

To grasp, catch, stack, or avoid objects, we need to infer their physical properties such as elasticity, mass, compliance, or friction (1–7). In most cases, we see objects before we interact with them, making vision the primary source of information to perceive and predict the physical world. Still, researchers do not yet fully understand the cues and computations the brain relies on to estimate the internal properties of objects (8–22). Unlike an object’s shape, size or identity, physical properties like mass or elasticity can only be *inferred* from the observed behavior of the object or substance (8–10,13,14,17), e.g., how a fluid flows, jelly wobbles or a ball bounces.

Consider a bouncing elastic object: How it bounces depends on many factors besides its elasticity, e.g., the initial direction and force with which it was thrown. An individual object can produce an infinite variety of trajectories, i.e., spatiotemporal paths, while objects with different elasticities can trace very similar paths depending on other factors, such as the object’s initial speed, height or direction of motion (**Figure 1A**). In previous work(8), we have shown that observers estimate the elasticity of bouncing cubes based on their motion trajectory. But, if there is no unique mapping between an object’s elasticity and its trajectory, how does the brain estimate the former from the latter?

**Figure 1.**
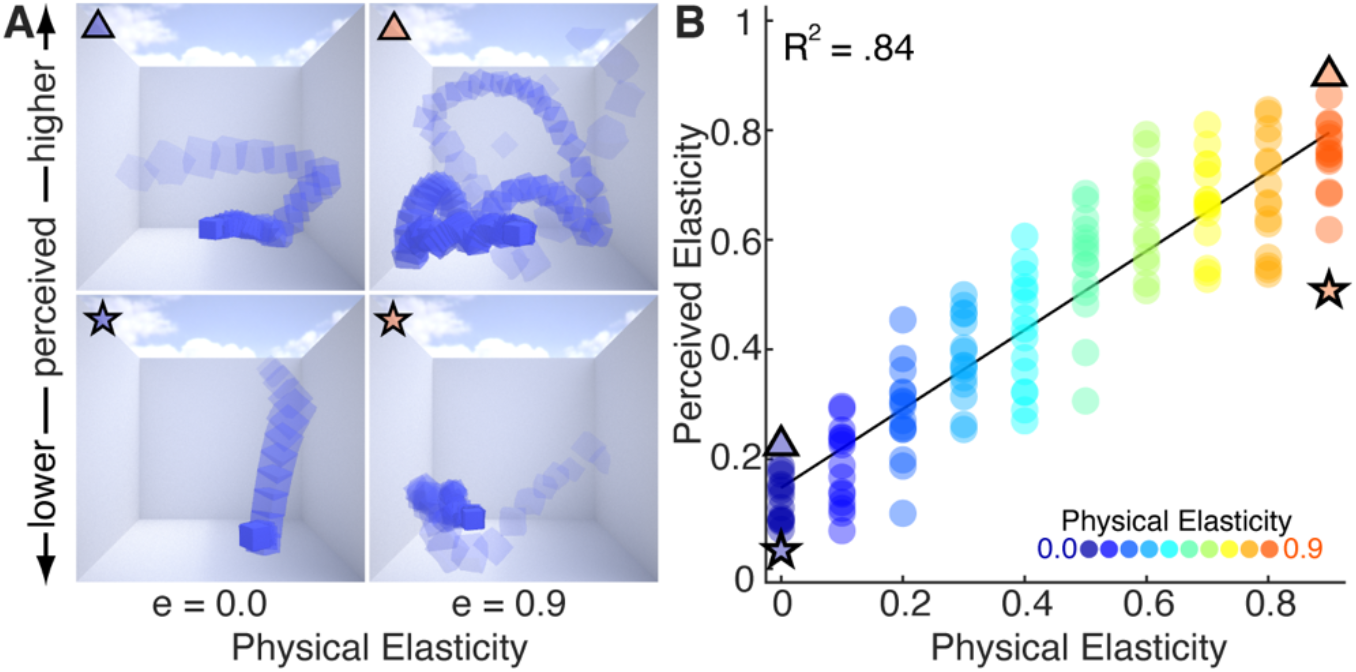
Stimuli and results of Expt. 1. **A)** Example stimuli of lowest (e = 0.0) and highest elasticity (e = 0.9), with animation frames overlaid for illustration purposes (see Video S1). Although both images in each row show the same cube (i.e., same elasticity), trajectories are different as we randomly varied initial speed, position, and orientation. **B)** Average elasticity ratings of Expt. 1 together with a linear fit. Dots of the same color show simulations of the same elasticity but varying initial parameters.

In their seminal work, Warren and colleagues (28) studied the perception of elasticity using simulations of idealized 2D elastic balls dropping perfectly vertically from five different heights. They found that under these conditions, observers use the relative height around a bounce (i.e. the *ratio of initial and final height*) to visually judge elasticity, and the *duration between two bounces* when the ball’s height is occluded. While their study elegantly isolated different cues and demonstrated that observers are sensitive to them, we were curious whether people use the same cues to judge elasticity in settings with more complex trajectories and when no single cue is a perfect determinant of elasticity (such as relative bounce height was in their study). In particular, while ideal spheres with zero initial velocity or spin produce perfect 1D spatial trajectories, even small increases in complexity can yield much more complicated behaviors for which it is unclear whether the same cues apply.

To test this, we investigated the perception of elasticity using simulations of bouncing cubes with randomly varying initial position, velocity and orientation. Under these conditions, the cubes traverse intricate 3D spatial trajectories including bouncing, tumbling and sliding. Importantly, this means that cues like the ratio of bounce heights or the duration of individual bounces are no longer uniquely mapped to the object’s elasticity. Instead, these—and other properties of the highly variable trajectories—are only statistically related to elasticity, suggesting that the visual system might need to combine multiple cues to infer the cube’s intrinsic properties.

Our work addresses two key questions: (1) How do people visually infer elasticity in naturalistic scenes, where no single cue alone perfectly predicts elasticity? (2) How does the brain learn to visually infer elasticity without ever having access to the ground truth? We reasoned that although individual trajectories vary, motion trajectories of the same elasticity will somewhat resemble each other in terms of their overall characteristic motion features, e.g., bounce height, speed of velocity decay and trajectory length. While no individual feature is perfectly diagnostic of elasticity, variations across different trajectories are also not random because they result from lawful physical constraints. By observing a number of examples, the brain could learn the dominant feature dimensions along which bouncing objects vary and represent elastic objects within the space of these features. A given heuristic (such as the bounce height ratio suggested by Warren and others) could be thought of as a special instance of this, in which the brain might identify just *one* feature along which trajectories are varying and thus elasticity judgments will rely on. However, another possibility is that the brain encodes elastic objects along multiple different features which would lead to a more robust representation in naturalistic settings. By considering different visual features, e.g., number of bounces and bounce height ratio, the brain could overcome the potential pitfalls of individual heuristics considered in isolation.

This idea leads to several testable predictions, which we evaluate here. First, motion features can be used to disentangle physical elasticity from other confounding factors (such as initial speed). Second, the relation between physical elasticity and motion features can be learned through observation alone. Third, either a single motion feature (i.e., a heuristic) or a robust combination of several features can explain the pattern of successes and failures in human perception. To test these assumptions, we employed a data-driven approach. For this purpose, we simulated 100,000 short (4 sec) trajectories of a bouncing cube in a room (**Figure 1A**). The cube’s elasticity (coefficient of restitution) varied from 0.0 (not elastic) to 0.9 (very elastic) in ten steps. Importantly, we also varied the initial position, orientation, and velocity of the cubes to gain 10,000 different trajectories for each level of elasticity. We validated that the simulations reproduce several crucial physical behaviors of bouncing objects (8). Only through simulation can we generate sufficient number and diversity of trajectories to identify and evaluate statistical regularities. We chose nonrigid, i.e., deformable, cubes as stimuli, because they result in chaotic and highly variable trajectories while being feasible in terms of the parameters to create and analyze them. Thus, they are a useful starting point for investigating the cues and strategies observers use when asked to judge elasticity. Of course, many other factors, such as the size and shape of the objects, or the elasticity of the floor can also impact bouncing objects’ trajectories. Yet, the range of behaviors exhibited by cubes already represents a significant generalization beyond previous research, and qualitatively spans many of the typical motions (bouncing, tumbling, sliding) one would expect to occur with real-world objects.

Next, we identified 28 candidate 3D motion features (**Figure 2A-D, Table S1**) based on the physics of bouncing objects, and previously proposed cues (28,29). We then determined how they statistically relate to physical elasticity in our dataset und used PCA to determine the optimal feature combination to predict elasticity. Our analysis revealed several competing hypotheses of how humans visually judge elasticity using motion features. In a series of carefully designed experiments, we selected stimuli that systematically decouple these highly correlated alternative hypotheses from one another and from ground truth, allowing us to find the one that best predicts human perception on a stimulus-by-stimulus basis. To begin, we first established human accuracy and consistency in elasticity perception in a random subset of our dataset as a benchmark to test out models of perception against.

**Figure 2.**
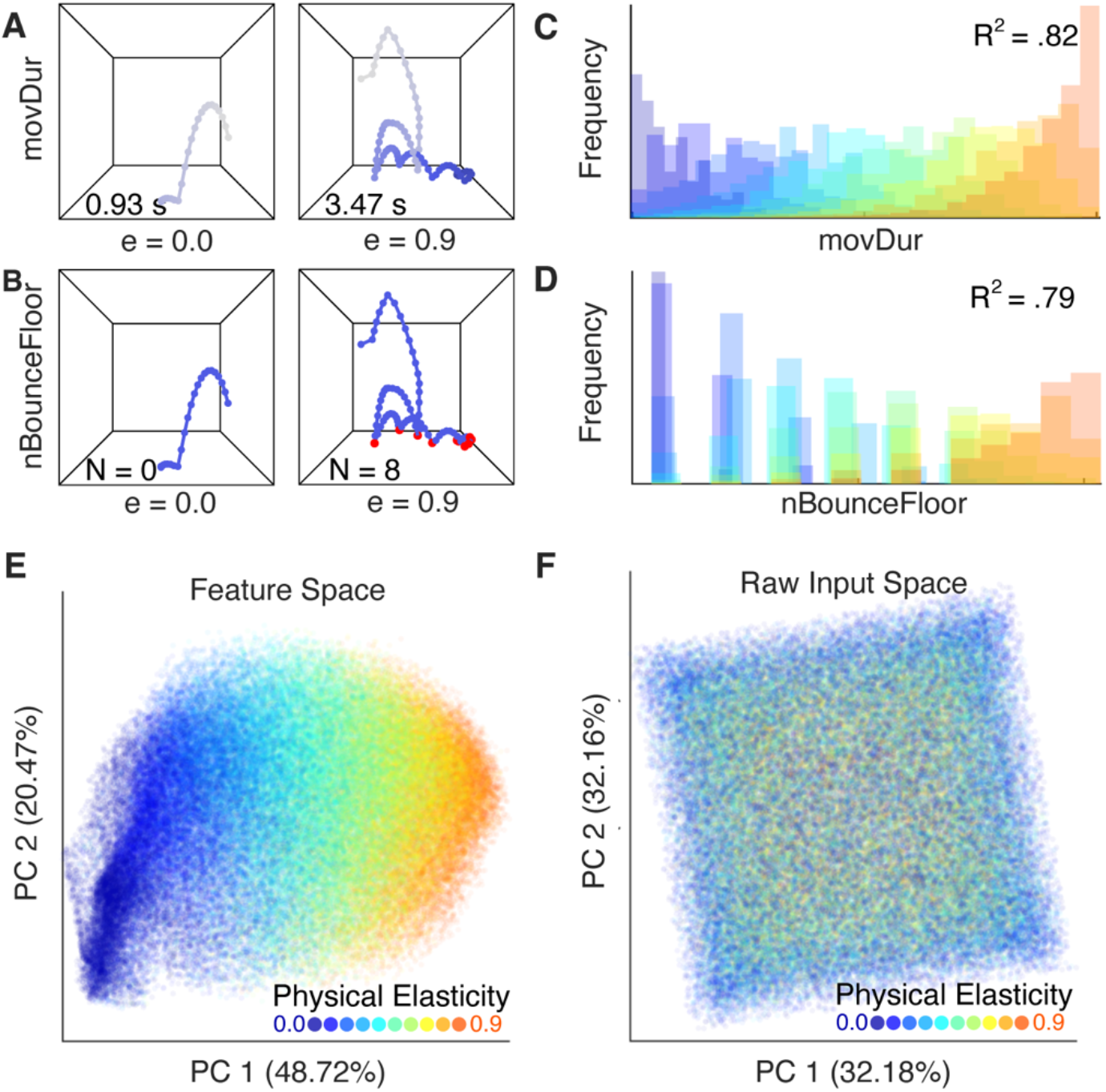
Spatiotemporal motion features of bouncing objects. **A)** Example trajectory of low (e=0.0) and high (e=0.9) elastic cubes, each dot represents one frame; color gradient represents movement duration. **B)** The same two trajectories, red dots represent bounces off the floor. **C)** Distribution of movement durations and **D)** distribution of “number of bounces off the floor” in the set of 100,000 stimuli, true elasticity is color-coded. **E)** All 100,000 simulations in the space of the first two PCs resulting from a PCA on the motion features (“feature space”). **F)** 100,000 trajectories in the space of the first two PCs resulting from a PCA on the raw trajectories. The PCs seem to be related to the position of the cube in 3D space, see **Figure S3E-F.** Note that although the initial position of the cube is uniformly sampled, its 3D position over time is biased due to gravity. This results in a tilted square in the 2D representation of the PCs.

## Results

### Observers accurately infer elasticity, but make systematic errors

Fifteen observers rated the apparent elasticity of bouncing cubes in 150 simulated animations—fifteen different trajectories for each of the ten elasticities (see **Methods** and **Figure 1A** and **Video S1**). Although the initial speed, position, and orientation of the cubes varied randomly, yielding widely variable trajectories, observers were very accurate at estimating the cube’s relative elasticity (**Figure 1B**). Average ratings increased systematically with physical elasticity (linear regression: R^2^ = .84, F(1, 148) = 748.73, *p* < .001). Yet, it is interesting to ask whether any of the residual variance was due to systematic deviations from veridicality rather than random noise. We found that the pattern of errors was not random but highly consistent between different observers (*r* = .91 ± .04; M ± SD) as well as within repeated ratings of the same individual (*r* = .90 ± .04). In fact, there was no significant difference between intra- and inter-observer variability (*t*(14) = 2.08, *p* = 0.056). What causes this systematic pattern of errors? The results give a first hint that when observers are asked to judge elasticity, they in fact latch onto particular characteristics of the bouncing motion. If humans represent elastic objects in terms of their characteristic motion features, perceptual errors should occur whenever a trajectory falls onto an “atypical location” in that feature space. We next sought to test this prediction by explicitly decoupling numerous motion cues from ground truth and one another. We held the physical elasticity constant yet observed large, systematic and predictable variations in elasticity judgments.

### Motion features disentangle physical elasticity from other latent factors

We propose that when asked to judge elasticity, observers rely on one or more features of the bouncing motion. To test this hypothesis, we explored a set of motion features derived from the cube’s 3D trajectory. We started with 28 potential features that between them capture many aspects of bounce trajectories (**Tables S1-2**; **Figures S1-2**). The features were selected by: (a) consideration of the physics of ideal bouncing objects, (b) proposals from previous literature (28,29), and (c) subjective observations of the simulations. Some features describe individual bounces (e.g., average bounce height, rebound velocity) or measure the coefficient of restitution in simplified, idealized settings (e.g., bounce height ratio). Others capture summary statistics that integrate over time and might be useful in realistic scenes that deviate from ideal conditions (e.g., number of bounces, movement duration; **Figure 2A-B**). Such statistics provide several different ways of measuring how quickly the object dissipates kinetic energy as it bounces around. All motion features are computable from observable quantities, i.e., positions and changes of positions over time (see **Methods**). Although object rotation and deformation are important for a complete physical representation of the object’s motion, we do not consider them here, as our previous findings show they have negligible effects on perceived elasticity in these stimuli (8). With this exception we aimed to achieve a comprehensive characterization of the trajectories by defining a diverse set of features to follow a data-driven approach and constrain our hypothesis space based on the data rather than a priori assumptions.

First, we evaluated how well each individual feature captured the variance across different elasticities. We found that many features varied systematically with physical elasticity (**Figure 2C-D & 3B, Table S1**). The most diagnostic features (which share the most variation with physical elasticity) were those that integrate information over time, such as movement duration or the number of bounces. Interestingly, we found that the two cues Warren et al (28) identified as important in judgments of elasticity from balls bouncing on 1D trajectories do not generalize to the more complex conditions we consider here. Specifically, *mBounceHtRatio* and *mBounceDur* share only 23.42% and 0.77% of the variance with physical elasticity, so they represent relatively poor elasticity cues for bouncing cubes—whereas they are perfectly correlated with elasticity for the special case considered by Warren et al (28).

We narrowed our hypothesis space by excluding features that explained < 5 % of the variance from further analysis. We found that the remaining 23 features were significantly correlated with one another across the set of 100,000 trajectories (mean absolute correlation, M = 0.48; see **Figure S3A-B**). To identify independent dimensions of variation, we applied principal component analysis (PCA) to the normalized and equalized motion features of all trajectories. Representing the trajectories in the space of the first two PCs reveals that physical elasticity varies largely along the first dimension (**Figure 2E**). Indeed, we found that ground truth elasticity and the first PC share 82.83% of their variance. In other words, physical elasticity emerges as the latent variable driving most variance in the feature representation of all trajectories. Although adding further PCs necessarily increases the explained variance of the dataset (**Figure S3C**), adding more PCs to a multiple linear regression model fitted to physical elasticity does not increase the shared variance by much (with all PCs: 86.25%). Moreover, while PC1 robustly predicts physical elasticity, it is mostly independent of the other latent parameters we used to initialize our simulations (e.g., velocity; all < 1.0%, **Figure S3D**). Thus, this linear combination of motion features (see **Figure S4** for feature loadings) successfully disentangles physical elasticity from other scene factors in our dataset. Notably, this feature weighting is not the result of a fitting process but emerges naturally and without supervision from the statistics across many examples. This underlines the potential of motion features to form a statistical appearance model of bouncing objects in a completely data-driven fashion. In the following, we test whether PC1 can explain the perceptual patterns found in Expt. 1 (‘multi-feature model’). Importantly, applying a PCA to the raw motion trajectories (**Figure 2F**) does not yield comparable elasticity estimates—highlighting the crucial role of appearance features.

### Optimal motion features predict elasticity perception

Next, we tested whether observers rely on a single motion feature (a heuristic) when estimating elasticity or instead combine features to a potentially more robust estimate, similar to PC1. Note that besides PC1, we also tested a more traditional precision-based maximum-likelihood estimation (MLE) model of cue combination in which the features are weighted according to their reliability, i.e., their weights are inversely proportional to their variance. The MLE model was overall highly correlated with the PC1 model (*r* = .98 over 100k simulations) but performed slightly worse in predicting elasticity (physical and perceived, see **Figure S5**). We therefore considered PC1 as the stronger cue combination model to test against individual heuristics.

Interestingly, we found that motion features that turned out to be diagnostic heuristics of *physical* elasticity, were also the best to predict perceived elasticity in Expt. 1 (**Figure 3B**). Strikingly, movement duration, the best feature for predicting *physical* elasticity, was also the best to predict *perceived* elasticity (R^2^ = .91, F(1, 148) = 1515.1, p < .001, **Figure 3A-B**). On a stimulus-by-stimulus basis, movement duration was a better predictor of human ratings than physical elasticity (evidence ratio (ER): w_movDur_/w_Physics_ = 1.51e+20). Can a combination of features outperform this? We find that a multi-feature model, i.e., PC1, is a very good predictor of perceived elasticity in Expt. 1 (linear regression: R^2^ = .89, F(1, 148) = 1210.5, p < .001, see **Figure 3B-C**). This is impressive given that the feature weighting was derived from observing the covariation of features in a large data set rather than a fitting procedure to the perceptual (or any) data. PC1 predicts perception better than the ground truth does (ER: *w*_FeatureModel_/*w*_Physics_ = 3.38e+13), but worse than movement duration (ER: *w*_movDur_/*w*_FeatureModel_ = 4.46e+06; *w*_movDur_ ≈ 1). However, the predictions of both models are strongly correlated (*r* = .95, *p* < .001, in 100k stimuli). In Expt. 2 we therefore systematically decouple their predictions.

**Figure 3.**
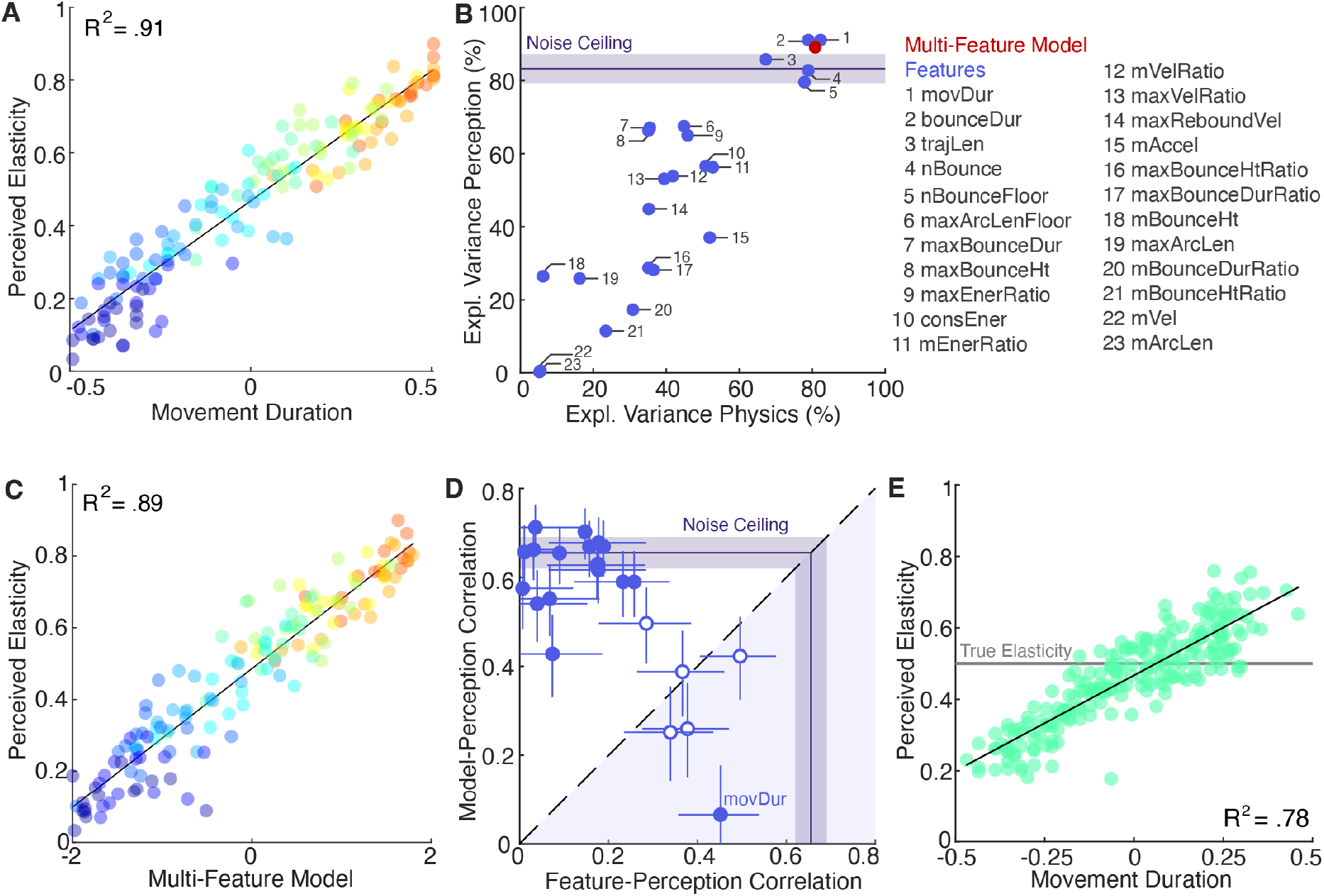
Prediction of perceived elasticity in Expts. 1 and 2 by competing models. **A)** Perceived elasticity (Exp. 1) as a function of the prediction made by the best single feature: movement duration. **B)** Explained variance in terms of perceived elasticity (Exp. 1; y-axis) and physical elasticity (x-axis) for individual features (blue) and the multi-feature model (PC1, red) in 100k stimuli. Noise ceiling shows the average explained variance between individual subjects and the average subject (± 95%-CI). **C)** Perceived elasticity (Exp. 1) as a function of the prediction made by the multi-feature model (PC1). **D)** Correlation of the pooled perceptual ratings (Exp. 2) with the multi-feature model (y-axis) and the individual features (x-axis). Each dot represents the correlations for one set of stimuli that were specifically selected to decouple the prediction of one feature from the model. Filled dots indicate a significant difference between the two correlation coefficients. Error bars show 95% confidence intervals. The noise ceiling shows the mean (± 95 % - CI) correlation between pooled responses and the average across features. **E)** Average elasticity ratings for all stimuli of Expt. 2 as a function of movement duration together with a linear fit. All stimuli had the same physical elasticity of 0.5 (grey line). Thus, all perceived differences in elasticity between stimuli are illusory. See **Video S2** and **Figure S6**.

### When observing complete motion trajectories people use movement duration as a heuristic to elasticity

The aim of Expt. 2 was threefold: First, we systematically decoupled the predictions of the multi-feature model from those of the movement duration heuristic to bring both models into conflict. Second, in order to test whether any of the other features are—individually—a better predictor of perceived elasticity, we systematically decoupled all other features from the multi-feature model. Since it is impossible to isolate each of the 23 features from all other features one by one, we decoupled each feature from the weighted combination of all features to test its causal contribution to elasticity perception. In doing so, we are able to overcome the purely correlational analysis reported so far and experimentally test 24 competing hypotheses at once, thereby going beyond previous studies (9–15). Third, because any good model of elasticity perception should be able to predict the pattern of errors on a stimulus-by-stimulus basis, all stimuli in this experiment had the same physical elasticity, i.e., all perceptual differences are illusory. This provides an even more stringent test of our 24 competing models, and a crucial high-level test of whether observers rely on cues and heuristics rather than an accurate internal model elasticity.

For this purpose, we simulated another 90,000 motion trajectories of the cube with medium elasticity (e = 0.5). From the total of 100,000 simulations of medium elasticity, we selected 23 sets of stimuli (one for each of the candidate motion features) for which individual feature and multi-feature model predictions were essentially uncorrelated (|*r*| < .05; see **Methods** and **Figure S6**). A new group of 30 participants judged the elasticity of these stimuli. Note that this rigorous stimulus selection process risks diminishing the very effects we seek to find: We first narrow the range of features by keeping elasticity constant and then select stimuli that, by definition, include outliers with a low correlation between a given feature and PC1.

Although these careful steps may have limited our statistical power, Expt. 2 provided clear results. For each feature, **Figure 3D** shows the correlation of perceived elasticity in the specific stimulus set (chosen for that feature) with the feature prediction (x-axis) and the multi-feature model prediction (y-axis). Seventeen features show a significantly lower correlation with perception than the multi-feature model (*p* < .0022, Bonferroni corrected). Only for one feature—movement duration—does the correlation with perception (*r* = .45) significantly exceed the multi-feature model (*r* = .07, *p* < .0022). In other words, when brought directly into conflict, movement duration can explain perceived elasticity better than a weighted feature combination. Thus, the high correlation between the multi-feature model and perception in Expt. 1 is presumably mediated by the contribution of movement duration (which has the third highest loading to PC1). Is movement duration also driving the high correlations between the multi-feature model and perception in the other stimulus sets of Expt. 2? **Figure S6E** shows the partial correlations between perception and single features vs. perception and multi-feature model prediction when controlling for the effect of movement duration. The correlations between perception and multi-feature model (*r* = .56 ± .14) decrease significantly when controlling for movement duration (*r* = .12 ± .11; *t*(21) = 12.97, *p* < .001), indicating that movement duration is indeed the driving factor. Across all stimuli, movement duration was—again—the best predictor of perceived elasticity (R^2^ = .78, F(1, 223) = 787.61, *p* < .001, see **Figure 3E and S5D**). Thus, the longer an object moved in the scene, the more elastic it appeared. Expt. 2 showed that this relation holds true even if physical elasticity is constant, leading to a powerful perceptual illusion. **Video S2** demonstrates these large, systematic, and robust illusory differences in apparent elasticity.

### Observers flexibly switch to another heuristic when movement duration is unobservable

Our everyday experience suggests that we are able to judge an object’s elasticity even without observing for how long the object moves, e.g., if someone catches it before it comes to rest. To study systematically how well people can estimate elasticity when this one cue is not available, we truncated a subset of the videos from Expt. 1 to exactly 1 second and presented these to a new group of 15 observers in **Expt. 3**, see **Video S3**. In these videos it was not possible to observe movement duration. Yet, we found that the average elasticity ratings increased systematically with physical elasticity (linear regression: R^2^ = .73, F(1, 78) = 215.41, *p* < .001, see **Figure S7A**) and showed a near-perfect correlation (*r* = .97, *p* < .001) with ratings for the full movies (Exp. 1; see **Figure 4A**). How do observers infer elasticity when movement duration cannot be observed? Do they rely on a different heuristic?

**Figure 4.**
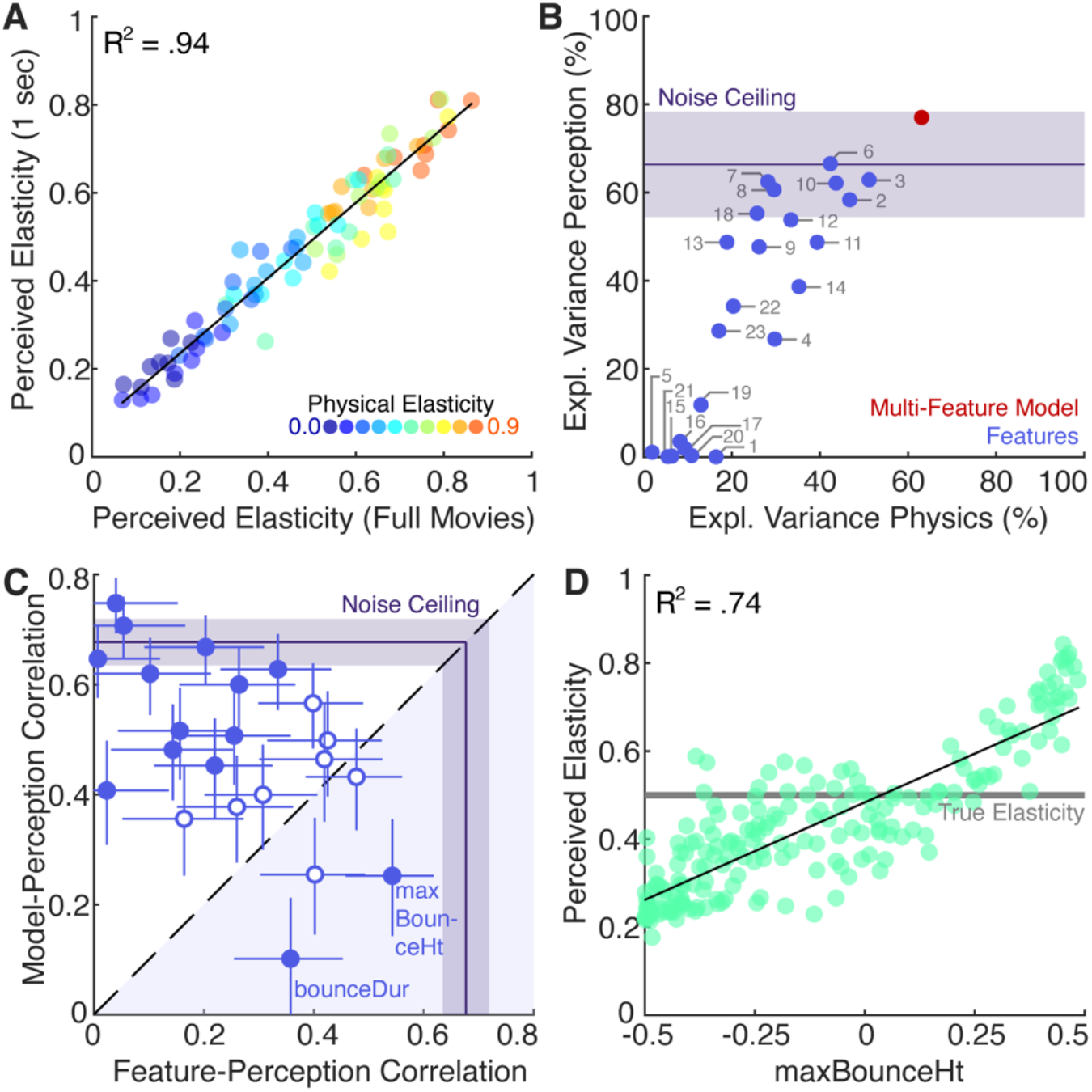
Results of Expts. 3 and 4 with truncated movies. **A**) Average perceived elasticity in 1-sec movie clips (Exp. 3) and full-length versions of the movies (Exp. 1). **B**) Explained variance in terms of perceived elasticity (Exp. 3; y-axis) and physical elasticity in 1-sec movies (100,000 stimuli). For a legend of individual features see Figure 2G. Noise ceiling shows average explained variance between individual subjects and the average subject (± 95%-CI). **C)** Correlation of the pooled perceptual ratings (Exp. 4) with the multi-feature model (y-axis) and the individual features (x-axis). Each dot represents the correlations for one set of stimuli, specifically selected to decouple the prediction of one feature from the model. Filled dots indicate a significant difference between the two correlation coefficients. Error bars show 95% CI. Noise ceiling shows the mean (± 95 % - CI) correlation between pooled responses and average across features. **D)** Average elasticity ratings (Exp. 4) as a function of the maximum bounce height together with a linear fit. See **Videos S3** and **S4**, and **Figure S7**.

Truncating the videos altered most feature values, not just movement duration. **Figure 4B** shows how well individual and combined features predict physical and perceived elasticity in 1-sec movies. The multi-feature model was the best at explaining both physics and perception and again better explains perception than ground truth physics (R^2^ = .77, F(1, 78) = 260.27, *p* < .001; ER: *w*_FeatureModel_/*w*_Physics_ = 296.05; see **Figure S7B**). Several individual features, particularly those measuring the presence of large bounces in the trajectory, also capture a large proportion of the variance in perceived elasticity. To disentangle these competing, but correlated hypotheses, we conducted **Expt. 4** following the same logic as in Expt. 2: From the dataset of 10^5^ medium elasticity cubes, we first identified simulations that had a movement duration of at least 1 sec. For this subset, we calculated the motion features for the first second and then selected 22 sets of stimuli (one for each feature except movement duration) in which the prediction of that feature individually was uncorrelated with the prediction of the multi-feature model. A new group of 30 observers estimated elasticity in these 1-sec stimuli.

Again, we found that one of the most diagnostic features—maximum bounce height—showed a significantly higher correlation with perception than the multi-feature model (*r* = .54 > *r* = .25, *p* < .0023, **see Figure 4C**) when brought directly into conflict, and that was the best predictor of perceived elasticity across all stimuli in Expt. 4 (R^2^ = .74, F(1, 197) = 565.56, *p* < .001, see **Figure 4D** and **S6C**). Thus, the higher the largest bounce was, the more elastic the cube appeared even if the true elasticity was equal (see **Video S4**). There was only one other feature—bounce duration—for which the correlation between feature and perception was larger than the correlation between multi-feature model and perception (*r* = .36 > *r* = .10, *p* < .0023). However, bounce duration did not vary much in the stimulus set, because in most simulations the cube would have bounced for longer than 1 second had the movie not been truncated (see **Figure S7E**). Therefore, bounce duration was only a diagnostic feature when it was notably shorter than one second. For most (12/22) features, we found that the multi-feature model predicted the data better than the individual features (*p* < .0023, Bonferroni corrected). Akin to the results of Expt. 2, these high correlations seemed to be driven by the best single feature, maximum bounce height (see **Figure S7D**). More precisely, the correlations between perception and multi-feature model (*r* = .49 ± .16 (M ± SD)) decreased significantly when controlling for maximum bounce height (*r* = .23 ± .09; *t*(21) = 6.65, *p* < .001).

In sum, Expt. 4 showed that observers reported robust perceptual differences between truncated stimuli even though all had the same physical elasticity. Perceived elasticity was best explained by one of the most predictive features, maximum bounce height. Intuitively this makes sense, as the maximal bounce height is easy to compute (i.e., requires only one position) and it occurs within the first second in most trajectories (94.1%, see **Figure S7F**). Taken together, this suggests that if unable to fully observe an object’s movement until it comes to a standstill, we instead form an impression of its elasticity based on the highest of the bounces that it makes.

## Discussion

Here we propose that when visually judging physical object properties, people often represent them in terms of their typical appearance—i.e., typical mid-level spatiotemporal features. Specifically, our results suggest that when asked to judge the elasticity of a bouncing object, observers judge how long the object moves. If the motion duration is cut short and cannot be observed, observers instead rely on the maximal bounce height to judge elasticity. Our findings can be contrasted with previous claims—based on 1D trajectories of bouncing balls (28)—that the visual system primarily uses bounce height ratios or bounce durations. These cues do not seem to generalize to the more complex and realistic conditions we consider here, as we find they are poor predictors of both physical elasticity and human judgments (**Figure 3B**). Instead, observers focus on either the overall motion duration or maximum bounce height. This implies a flexible and computationally efficient strategy.

While this study is not the first to suggest a role of mid-level features in the estimation of physical properties (9–15,17,30), it overcomes three limitations of previous work. First, we evaluate a diverse set of potential cues in a large dataset and thereby show how—in principle—observing the variations of motion features in many examples spontaneously reveal elasticity and establish which features (or their combination) are best at doing so. Second, to the best of our knowledge, no study has yet *manipulated* the proposed visual cues to physical properties in naturalistic stimuli. Here, we achieved such manipulation by using a large dataset to identify stimuli that decouple the inherently correlated predictions of different models. Third, we identified *illusory* stimuli, i.e., movies with the same physical elasticity but different values of various cues, in which observers substantially and consistently misperceived elasticity. Thus, we can predict specific perceptual errors on a stimulus-by-stimulus basis. Our findings have implications on both theoretical and methodological levels.

### Are perceived differences of simulated elasticity “illusions”?

When two objects with the same physical parameters are systematically and predictably perceived as different, this matches the common definition of an “illusion”. In the Müller-Lyer illusion (54), two lines that have physically the same length are seen as different. In the checker-shadow illusion (55, 56), two patches that have physically the same luminance are seen as different. Here, we use the term “illusion” to refer to an equivalent situation in which the simulated physical elasticity is the same for two stimuli, but elasticity ratings are substantially different. We suggest that this is not contradicted by the fact that this inference can be considered reasonable. Even ideal observers make erroneous inferences when the image information is deceptive (57). For example, the Ponzo illusion is still generally called an “illusion” even though it is reasonable for the visual system to infer different line lengths assuming the lines are at different distances from the observer. The same is true for elasticity estimates from motion features.

### Learning

We have previously hypothesized that by observing the outside world and its inherent statistical relations (31,32), the brain can learn many dimensions along which objects in our environment vary—in an unsupervised manner. The statistical appearance model proposed here is not intended as a model of this learning process, but rather a proof of principle about the learnability of the cues and their role in perception. We found that by observing various motion features of bouncing cubes, elasticity emerges as the main dimension of variation in our multi-feature space. The motion features themselves were explicit operationalizations of our hypotheses and not the result of learning. This approach allowed testing the contribution of a large, yet testable number of interpretable features and their combination. Would similar features emerge from unsupervised or self-supervised learning algorithms? It would be interesting to investigate this question within different frameworks, from deep learning to program learning or simulation-based learning. However, it would be practically impossible to test the individual contribution of the thousands of features in the trained network to perceived elasticity. Yet, here, it is precisely this decoupling of hypotheses that enabled us to predict human perception on a stimulus-by-stimulus basis.

### Mid-level features

One of our key findings is that when asked to estimate the elasticity of bouncing objects, observers judge the movement duration or the maximal bounce height in case the duration is visibly cut short. Crucially, our findings also imply that the brain does represent *multiple* features of bouncing objects at a time but does not combine them in the sense of classic cue combination (33) to estimate the latent parameter. Our results suggest, that ‘estimating elasticity’ means determining the relative position of the observed object on the feature manifold.

Why would the brain rely on these and not on other features? Presumably, the most effective features are (1) strongly related to elasticity, (2) vary in salient ways across stimuli and (3) inexpensive to compute. Movement duration and maximum bounce height are both strongly related to elasticity and capture important events in the observed motion, i.e., the largest bounce and the end of the motion. While it is not trivial to determine the computational costs of different features, it seems plausible to assume that single measures, e.g., height, will be cheaper than their derivatives or ratios. In that sense, both features are among the computationally simplest we tested. They are also quantities the visual system estimates reliably and accurately (34–37).

A key assumption of our model is that the observed motion is due to a bouncing elastic object, as opposed to some other cause (e.g., animate motion (38), fluid flow (13,14,39)). If applied to other trajectories, e.g., a feather gliding in the wind, the resulting ‘elasticity estimate’ would be meaningless. Future research needs to investigate whether the other features help identify the stimulus as a bouncing object in the first place.

### Image-computability and modelling visual processes

The motion features we tested here are *stimulus-computable*, i.e., they can be computed from changes in position of the surface points over time. This allows us to assume ideal conditions to set upper bounds on the utility of specific cues. Thus, when our experimental data rules out a particular cue, we can be sure it is not due solely to limits of our modelling or the particular implementation details of how the cue is computed from the raw sensory data. However, because the features are computed using a perfect representation of the object’s trajectory, they oversimplify the input available to biological brains. For example, humans have a more accurate representation of image-plane motions than motion in depth (40,41), and may not be equally sensitive to all velocities in these displays. To transform the heuristic model into a truly image-computable one, future work will need to incorporate aspects of low-level vision, including object segmentation and depth perception. Yet, we reasoned that important insights into the estimation of material properties can still be gained even without fully modeling all preceding processing stages. Indeed, given that our experimental findings rule out most of the hypothesized cues, future work should focus on modelling how the visual system derives motion duration and maximum bounce height from responses of motion-selective units in V1, MT and other cortical regions. Another interesting novel challenge will be to model the processes by which the visual system switches between cues.

### Generalization

One key limitation of our study was that we held certain factors constant that would likely vary across scenes in the real world: size, shape, mass and friction of the bouncing object; elasticity and friction of the floor; and the layout of the surrounding surfaces, to name a few examples. It is possible that elasticity would not align so closely with the first principal component of the space if we had varied additional physical properties. Yet, deformable cubes do produce highly diverse and complex trajectories, featuring many of the typical behaviors we expect to observe with more varied conditions. They change directions in seemingly unpredictable ways, vary in bounce heights non-monotonically, and roll or slide along the floor. We have shown that certain visual motion features, particularly motion duration and maximum bounce height, robustly generalize across these conditions. Their robustness likely extends to even more varied conditions, where participants may rely on them even more heavily as heuristics—precisely because they are among a small set of features that remain stable and easily measurable across diverse scenarios. In an experimental setting, it would of course be possible to break the relation between motion features and elasticity. For example, if the floor was completely inelastic, like sand, no object would rebound. It is, however, unlikely that human observers would be able to estimate the objects’ elasticity in these cases. Thus, although motion features would not capture physical elasticity, they might still be reliable predictors of *perceived* elasticity. Because our model is stimulus computable, such hypotheses can be easily tested in future research.

### Simulation vs. heuristics

A current topic of debate is the extent to which physical perception and reasoning proceed through sophisticated but computationally costly internal simulations (19–22,42) or cheaper but potentially less accurate heuristics (9,10,43,44). How do our results fit into this theoretical spectrum?

Representing objects and materials in terms of their appearance features entails an understanding of the observable consequences of natural variations between objects, e.g., the ways in which elastic objects bounce. Yet, the resulting estimation strategy appears much more like a classic heuristic (e.g., “the longer it moves, the more elastic it is”), than the result of detailed internal simulations. For most practical purposes, the goal of the visual system is to anticipate what is going to happen next—whether an object will change direction, when it will come to rest, etc. Capturing the characteristics of the ‘bouncing motion’ of the object is perhaps more useful than inferring its intrinsic coefficient of restitution.

We suggest that our results also provide a hint at how the brain derives such heuristics from observation alone, and of how it switches from using one feature to another (i.e. when there is no variation along the first feature dimension). This does not mean that observers *cannot* simulate possible future behaviors of objects, such as how the trajectory of a bouncing cube continues, just that they may not choose to do so when simpler yet near-optimal heuristics are available. This interpretation is consistent with previous work by Battaglia and colleagues (20), who found that when a simple heuristic is a more efficient and optimal way to make a prediction, observers tend to use heuristics rather than simulation. Thus, we suggest that observers can draw on different forms of computation, but do so taking into consideration the relative costs and demands of the specific task at hand—an example of *bounded* or *computational rationality* (45,46). When asked to infer a single parameter (e.g., elasticity) from an observed trajectory, time- and energy-consuming simulations represent a poor allocation of resources when a simple read-out from the feature estimation provides high accuracy. However, visual features are likely too inaccurate when making time- or location-critical predictions about an object’s future trajectory (7,47,48). Under these conditions, the additional costs associated with internal simulation may pay off. Similarly, when no standard heuristics apply, observers may use simulation even for physical inference of material properties as shown by Hamrick et al (21). Future studies should further investigate the cognitive strategies humans employ across various circumstances and the metacognitive processes that govern them.

## Conclusion

Visually estimating physical object properties is a crucial, yet computationally challenging task. The visual input is highly ambiguous because an object’s behavior depends on numerous entangled factors. Estimating the elasticity of a bouncing object requires disentangling the different causal contributions of elasticity, initial speed, position, and other factors. Using a ‘big data’ approach, we showed that representing trajectories in terms of their characteristic spatiotemporal features—such as the maximum bounce height or movement duration—yields elasticity estimates that are inexpensive to compute and robust to external factors. Our experiments suggest that the brain estimates elasticity by flexibly switching between a few single-feature heuristics based the information available in the stimuli. Our model explains both the broad successes and the systematic failures of human elasticity perception and correctly predicts a novel illusion in which appearance features maximally diverge from ground truth. Observers can draw on multiple cues and computations, and appear to select strategies with lower computational costs.

## Methods

### Subjects

Ninety undergraduate students (68 females) from the University of Giessen participated in the experiments (15 in Exp. 1 and 3, 30 in Exp. 2 and 4). Their average age was 24 years (SD = 3.5). All participants were naïve with regard to the aims of the study and gave written informed consent before the experiment. Participants were compensated with 8€/h. The experimental procedure was in accordance with the declaration of Helsinki and approved by the local ethics committee (LEK FB06).

### Physical simulations

The dataset was created with the Caronte physics engine of RealFlow 2014 (V.8.1.2.0192; Next Limit Technologies, Madrid Spain), a 3D dynamic simulation software. The dataset contains 100,000 simulations of a cubical object (0.1 m^3^) bouncing in a cubical room (1.0 m^3^). We varied the cube’s elasticity (coefficient of restitution) in ten equal steps from 0.0 to 0.9. We created 10,000 simulations for each level of elasticity by randomly varying its initial velocity, orientation, and position. We simulated 121 frames at 30 fps of the cube moving under gravity. We simulated additional 90,000 trajectories of medium elasticity (0.5) by randomly varying initial velocity, orientation, and position and used the total 100k simulations of medium elasticity to select stimuli in Expts. 2 and 4. We provide 3D positions of the cube’s corner and center of mass (CoM) across all frames for all 190k simulations here: https://doi.org/10.5281/zenodo.17196171.

### Motion features and multi-feature model

We calculated 28 motion features based on the CoM and the eight corners of the cube for all 100,000 simulations. The end of the cube’s movement was defined as the point at which its velocity dropped below 0.003 m/s, since simulated velocity never reaches zero. All other features were computed only for the frames in which the cube was moving. The exact definition of all 28 motion features is described in **Table S1** and **Figures S1-2**. Next, we normalized every motion feature to a range between [0.0, 1.0] and equalized their histograms. We determined the R^2^-score, the shared variance with physical elasticity, for each feature and excluded features from further analysis if they explained <5% of the variance. We performed a principal component analysis (PCA) with the remaining 23 features. The resulting scores of the first principal component (PC1) were used to predict perceived elasticity, see **Figure S3**.

### Stimuli

Expt. 1 contained 15 stimuli per level of elasticity, randomly selected from the original dataset (i.e., 150 stimuli). For Expt. 2 we selected 225 stimuli that systematically decoupled the predictions of each individual feature from both the multi-feature model and physical elasticity. Specifically, for each of the 23 features we chose ten stimuli from the medium elasticity dataset for which the predictions of the individual feature and the multi-feature model were uncorrelated (|*r*| < 0.05), while spanning the widest possible range on both dimensions (**Figure S6A**). In Expt.s 1 and 2, each stimulus was presented for the full duration of the cube’s movement. In Expt.s 3 and 4, only the first second of each stimulus was presented (and no stimulus had a movement duration that was shorter than 1 sec). For Expt. 3, we used a subset of eight stimuli per elasticity level from the stimuli of Expt. 1 (i.e., 80 stimuli). For Expt. 4, we selected 213 stimuli that systematically decoupled the predictions of each individual feature (except movement duration) from both the prediction of the multi-feature model and physical elasticity. The selection procedure was the same as in Expt. 2, but all stimuli were truncated to exactly one second. The simulations selected as stimuli were rendered using RealFlow’s built-in Maxwell renderer and are available for download at https://doi.org/10.5281/zenodo.17196171.

### Set up

All experiments were conducted using the same setup. Stimuli were presented on an Eizo LCD monitor (ColorEdge CG277; resolution: 2560 × 1440 pixels; refresh rate: 60 Hz). Participants used a chin rest to maintain a constant viewing distance of 54 cm. At this distance, the stimuli had a visual angle of 19.6 × 19.6 degrees.

### Procedure

All experiments followed the same procedure. Participants were instructed to watch a short movie of an object and rate its elasticity. Elasticity was defined to them as the property that distinguishes for example a bouncy ball from a hacky sack. On each trial, one stimulus was presented repeatedly until a response was given. Participants adjusted a slider on a horizontal rating bar to indicate their rating between ‘not elastic’ to ‘very elastic’. All stimuli were presented in random order including three repetitions per stimulus. Before the main experiment, participants completed ten practice trials, one for each level of elasticity (unknown to participants) to provide an impression of the stimulus range without biasing their response scale. The experimental code was written in Matlab 2018a using Psychtoolbox 3 (49–51).

### Analysis

In all experiments, we averaged across repetitions to obtain one rating per stimulus from every participant. For Expt.s 1 and 3, we calculated the average across participants, as well as inter- and intra-observer variability (standard deviation). We fitted linear regression models to the average ratings using either physical elasticity, the multi-feature model, or each of the individual feature as predictors. Models were compared using AIC values, specifically their Akaike weights and evidence ratios (52). Akaike weights represent the probability that a given model is the best among those tested, while evidence ratios indicate the relative likelihood of two competing models given the data. For Expt.s 2 and 4, we pooled the data across participants. For each feature’s stimulus set, we computed correlations between pooled ratings and both the corresponding feature prediction and the multi-feature model prediction. Pooling (rather than averaging) allowed more reliable estimation of the correlation coefficients based on full trial counts rather than just 10 averages. For each feature, the resulting correlation coefficients were compared using a two-tailed significance test for dependent groups with one overlapping variable (53). Additionally, we computed the explained variance in perceived elasticity (averaged across participants) for each feature and the model across *all* stimuli, independent of the stimulus set.

## Ethics

The experimental procedure was in accordance with the declaration of Helsinki and approved by the local ethics committee at Giessen University (LEK FB06).

## Data Accessibility

All 190k simulations, experimental stimuli, data and code necessary to analyze the motion features and reproduce the results and figures presented in this manuscript are publicly available here: https://doi.org/10.5281/zenodo.17196171

## Declaration of AI use

We have not used AI-assisted technologies in creating this article.

## Author Contributions

VCP, FSB, and RWF conceived and designed the study, developed the features and computational model. VCP simulated the data set, rendered the stimuli, collected and analyzed the data, made the figures, and wrote a first draft of the manuscript. All authors discussed the results and wrote and approved the manuscript.

## Conflict of interest declaration

We declare we have no competing interests.

## Funding

This research was funded by the Deutsche Forschungsgemeinschaft (DFG, German Research Foundation; Project 222641018–SFB/TRR 135 TP C1 and Project PA 3723/1-1) and by the European Research Council (ERC) Advanced Grant “STUFF” (Project 101098225).

## Acknowledgments

The authors thank Britta Fritz, Saskia Honnefeller, Judith Kanehl, and Jasmin Kleis for their help with data collection.

## Notes

https://doi.org/10.5281/zenodo.17196171

### Competing Interest Statement

The authors have declared no competing interest.

### Summary of Updates

This is the peer-reviewed, revised and accepted version of our manuscript.

