## Supplemental Information for "Visual cues and strategies for perceiving elasticity"

Main article: https://doi.org/10.1098/rspb.2026.0483

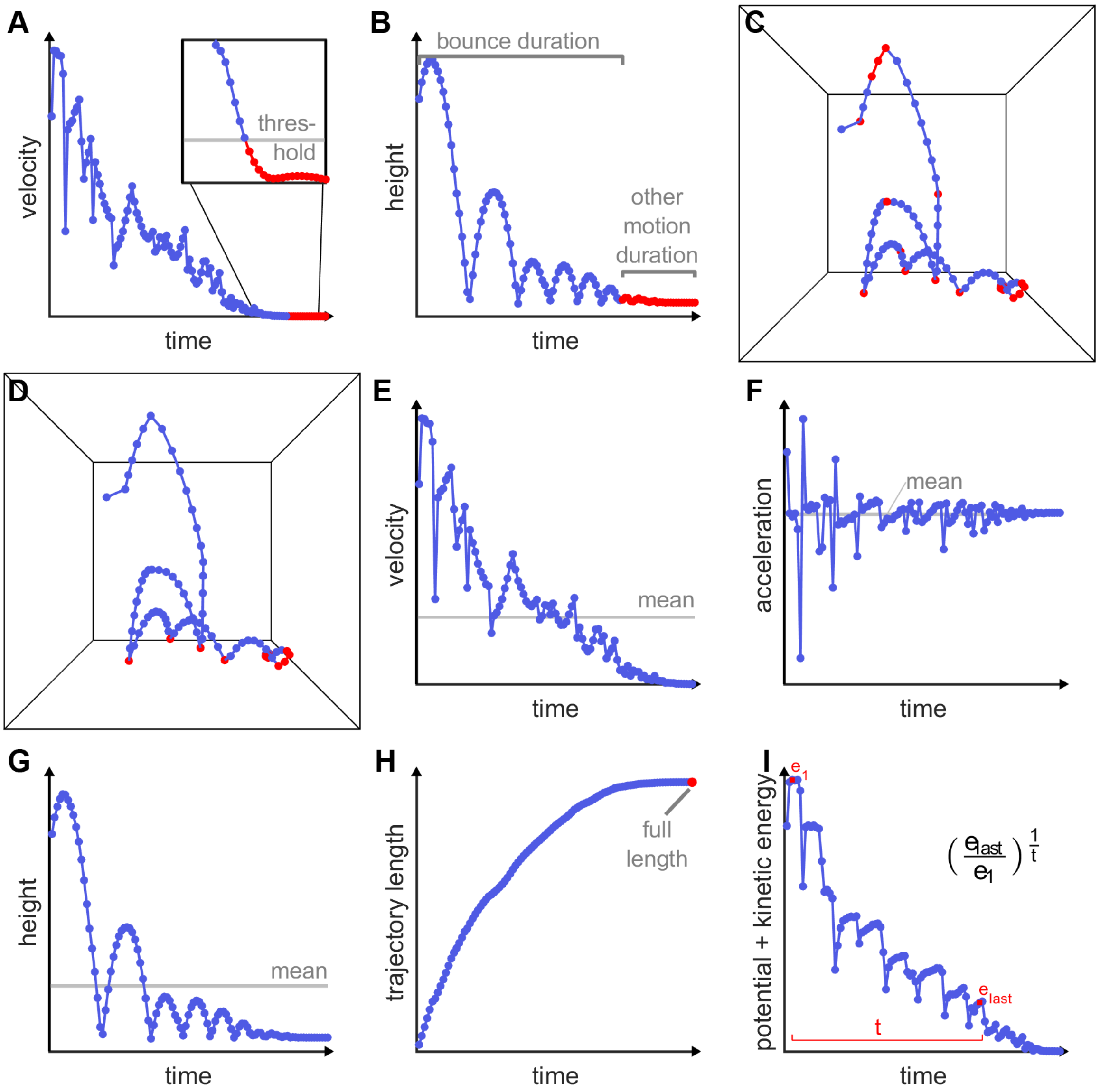


**Figure S1. Visualization of motion features integrating statistics over time. A) Movement duration:** Movement duration was determined using a velocity threshold. Plot shows the velocity of the CoM of an example trajectory as a function of time. Box shows zoom-in on data and threshold, with which we determined the end of the motion. A threshold was necessary because the simulations never truly reach a velocity of zero. **B) Bounce and other motion duration:** Plot shows the height of the CoM of an example trajectory as a function of time for the duration during which the cube was moving. The duration until the end of the last bounce (i.e. when the cube landed) was defined as *bounce duration* (blue). The remaining time during which the cube was moving (e.g. sliding, wobbling, rolling), but not bouncing, was defined as '*other motion duration*' (red). **C) Number of bounces:** Plot shows the 3D trajectory of the CoM of an example simulation; black lines outline the room. Each dot represents one frame; red dots represent bounces from any wall. A bounce was detected if one of the cube's corners or its CoM (± ½-edge length) was in contact with a wall, preceded and followed by all points of the cube being in the air. By this definition, a bounce can span several frames if one of its edges is in contact with the wall in subsequent frames (see three red dots showing a bounce from the ceiling). The position of the cube was interpolated between frames to detect bounces outside the frame rate. **D) Number of bounces from the floor:** Plot shows the 3D trajectory of the CoM of an example simulation; black lines outline the room. Each dot represents one frame; red dots represent bounces from the floor (same criteria as in C). **E) Mean velocity:** Plot shows the velocity of the CoM of an example trajectory as a function of time for the duration during which the cube was moving. Grey line indicates the mean across time, which was used as a feature. **F) Mean acceleration:** Plot shows the acceleration of the CoM of an example trajectory as a function of time for the duration during which the cube was moving. Grey line indicates the mean across time, which was used as a feature. **G) Mean height:** Plot shows the height of the CoM of an example trajectory as a function of time for the duration during which the cube was moving. Grey line indicates the mean across time which was used as a feature. **H) Trajectory length:** Plot shows the cumulative length of the trajectory of the CoM of an example trajectory as a function of time for the duration during which the cube was moving. Red dot indicates the full length of the trajectory, which was used as a feature. **I) Conserved energy:** Plot shows the sum of potential and kinetic energy of an example cube as a function of time for the duration during which the cube was moving. Energy was approximated as the potential maximal speed of fall, which is proportional to the sum of kinetic and potential energy. We calculated the energy *e* based on the velocity *v* and height *h* of the cube (i.e., ignoring its rotation and deformation): $e=v+\sqrt{2gh}$, where *g* denotes gravity (9.81 m/s2). Conserved energy was defined as the average energy during the last flight phase divided by the energy during the first flight phase (both in red) normalized by the time that has passed between the two time points.

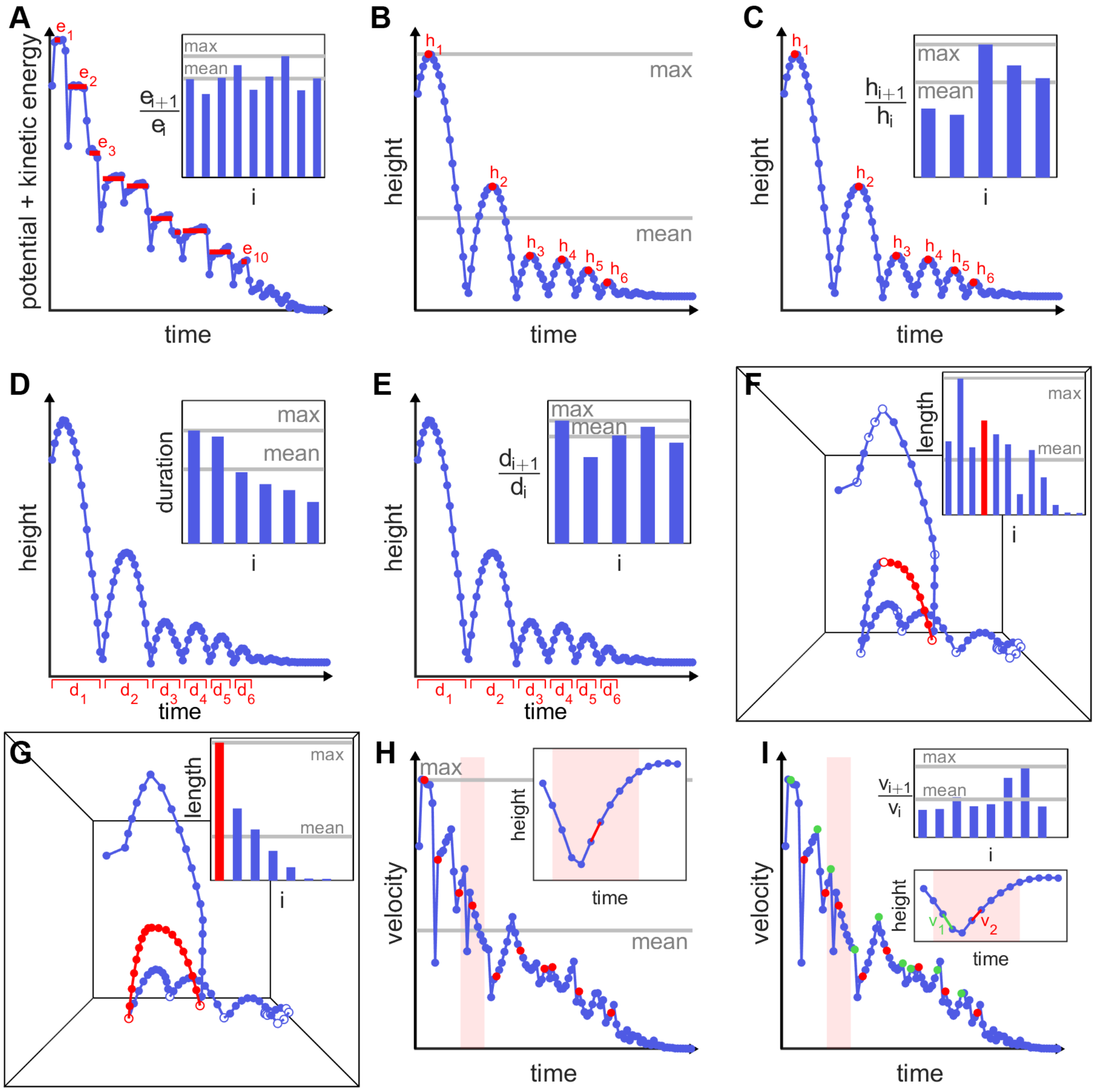

**Figure S2. Visualization of motion features characterizing bounce events. A) Mean and maximum of energy ratio per bounce:** Plot shows the sum of potential and kinetic energy of an example cube as a function of time for the duration during which the cube was moving. Red lines indicate the average energy during flight phases. The small plot shows the energy ratio for pairs of subsequent bounces, i.e. the ratio between the average energy in two subsequent flight phases. The mean and maximum of all energy ratios (grey lines) were used as features. **B) Mean and maximum bounce height:** Plot shows the height of the CoM of an example trajectory as a function of time for the duration during which the cube was moving. Red dots indicate the height of every bounce. Grey lines indicate the mean and maximum across time, which were used as features. **C) Mean and maximum bounce height ratio:** Plot shows the height of the CoM of an example trajectory as a function of time for the duration during which the cube was moving. Red dots indicate the height of every bounce. Small plot shows the height ratio for pairs of subsequent bounces. Grey lines indicate the mean and maximum across bounces, which were used as features. **D) Mean and maximum bounce duration:** Plot shows the height of the CoM of an example trajectory as a function of time for the duration during which the cube was moving. Red lines indicate the duration of every bounce. Small plot shows the duration of each bounce together with the mean and maximum across time (grey lines), which were used as features. **E) Mean and maximum bounce duration ratio:** Plot shows the height of the CoM of an example trajectory as a function of time for the duration during which the cube was moving. Red lines indicate the duration of every bounce. Small plot shows the duration ratio for pairs of subsequent bounces. Grey lines indicate the mean and maximum across bounces, which were used as features. **F) Mean and maximum length of bounce arcs:** Plot shows the 3D trajectory of the CoM of an example simulation within the room. Each dot represents one frame; white dots represent bounces from any wall. Red dots indicate frames of one example arc starting from the floor and ending at the front wall (difficult to see in the 2D projection). Small plot shows the arc length, i.e., the cumulative distance the cube traveled between two bounces, of all bounces (same example in red). Grey lines show the mean and maximum across all bounces, which we used as features. **G) Length of bounce arcs from floor:** Plot shows the 3D trajectory of the CoM of an example simulation within the room. Each dot represents one frame; white dots represent bounces from the floor. Red dots indicate frames of one example arc from floor to floor (ignoring the bounce from the front wall in between). The small plot shows the arc length, i.e. the cumulative distance the cube travelled between two bounces, of all bounces (same example in red). Grey lines show the mean and maximum across all bounces, which we used as features. **H) Mean and maximum rebound velocity:** Plot shows the velocity of the CoM of an example trajectory as a function of time for the duration during which the cube was moving. Red dots indicate the rebound velocity after a bounce. Grey line indicates the mean and maximum across time, which were used as features. Small plot shows the height of the CoM for few example frames highlighted in the red shaded area. The red segment corresponds to the rebound velocity shown in the main plot. **I) Mean and maximum velocity ratio:** Plot shows the velocity of the CoM of an example trajectory as a function of time for the duration during which the cube was moving. Red dots indicate the rebound velocity after a bounce. Green dots indicate the incident velocity. Lower small plot shows the height of the CoM for few example frames highlighted in the red shaded area with green and red corresponding to the incident and rebound velocities in the shaded area of the main plot. The velocity ratio was calculated by dividing the rebound by the incident velocity. The upper small plot shows the velocity ratio for every bounce. Grey line indicates the mean and maximum across time, which were used as features.

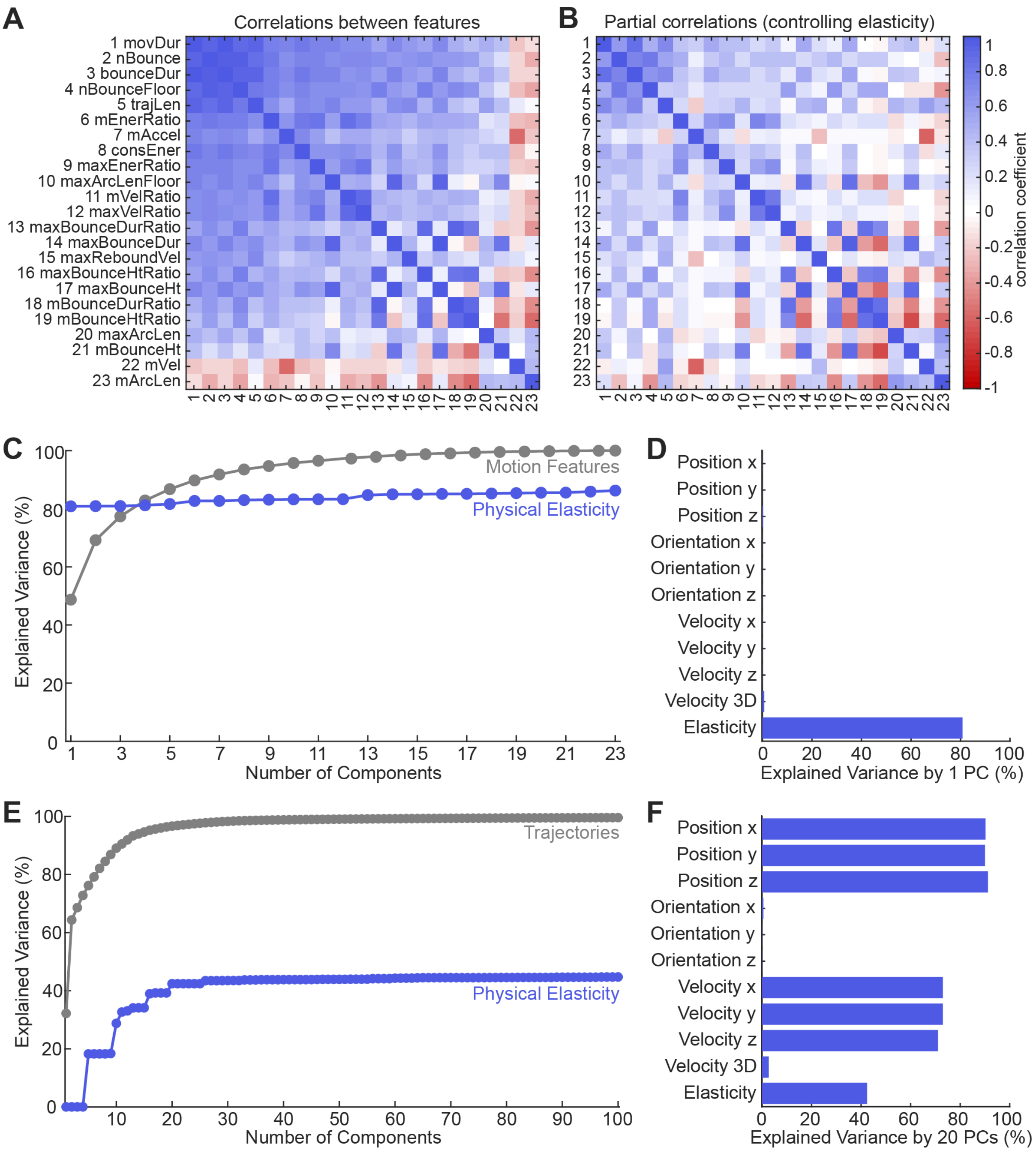


**Figure S3. Multicollinearity of motion features and PCA results, related to Figure 2. A)** The correlation matrix shows how much each feature correlates with every other feature. Motion features are ordered from top to bottom (and left to right) according to how much variance they explain in terms of physical elasticity. **B)** The correlation matrix shows the partial correlations between all motion features, controlling for the effect of physical elasticity. This analysis suggests that the multicollinearity (as seen in A) is partly because all features correlate strongly with physical elasticity. **C)** Cumulative explained variance through increasing numbers of PCs from the PCA calculated on the 23 motion features for the input data (i.e., the motion features; grey) and from a regression of an increasing number of resulting PCs to predict physical elasticity (blue). **D)** Explained variance of PC 1 (based on motion features) for all latent variables in our simulations. PC 1 predicts physical elasticity very well and is insensitive to variations in the initial position, orientation, and velocity. **E)** Cumulative explained variance through increasing numbers of PCs from the PCA calculated on the raw trajectories for the input data (i.e., the trajectories; grey) and from a regression of an increasing number of resulting PCs to predict physical elasticity (blue). The plot shows the first 100 of 1098 PCs. **F)** Explained variance of the first 20 PCs (of the PCA on raw trajectories, see E) as obtained by combining the 20 PCs optimally in a linear regression to predict each of the latent variables of our simulations.

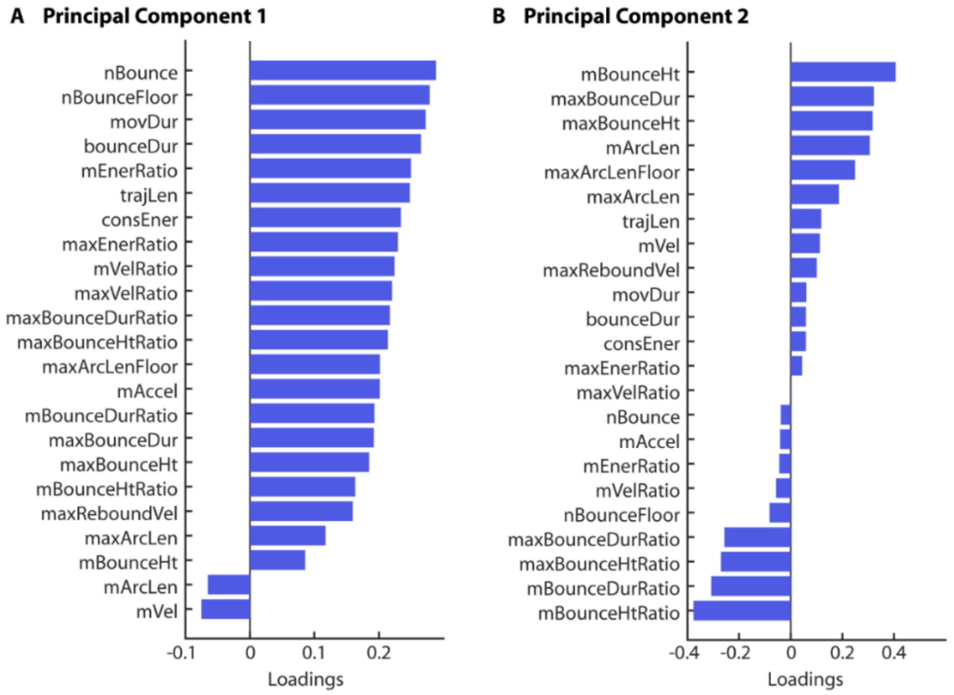


**Figure S4. Loadings of the first two principal components of the feature space, related to Figure 2. A**) Feature loadings of principal component 1 are dominated by motion features that integrate motion characteristics over time. **B**) Feature loadings of principal component 2 are dominated by motion features that characterize the height, duration and length of individual bounces.

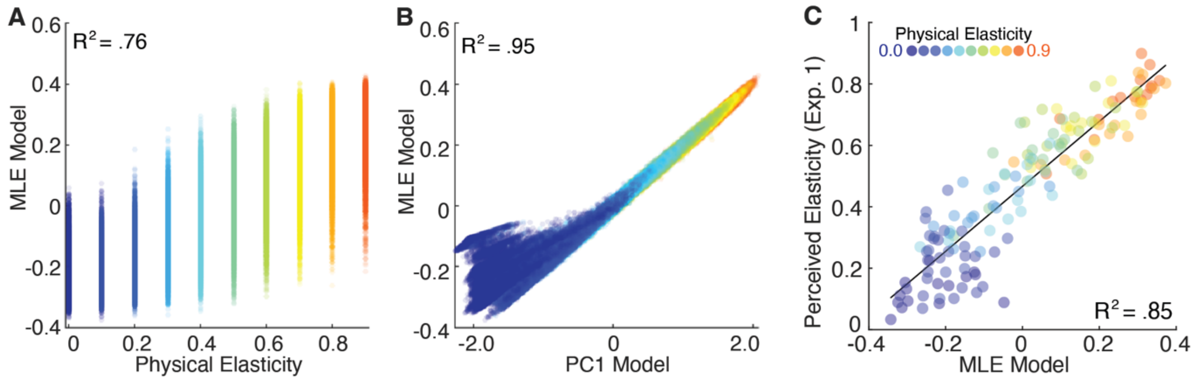


**Figure S5. Results of the Maximum-Likelihood (MLE) Model. A)** MLE Model prediction for 100,000 simulations as a function of their physical elasticity (color-coded). Dots of the same color show simulations of the same elasticity but varying initial parameters. **B)** MLE Model prediction for 100,000 simulations as a function of the prediction of a weighted feature combination as in PC1. Dots of the same color show simulations of the same elasticity but varying initial parameters. **C)** Average elasticity ratings from Experiment 1 as a function of MLE Model prediction together with a linear fit. Dots of the same color show simulations of the same elasticity but varying initial parameters.

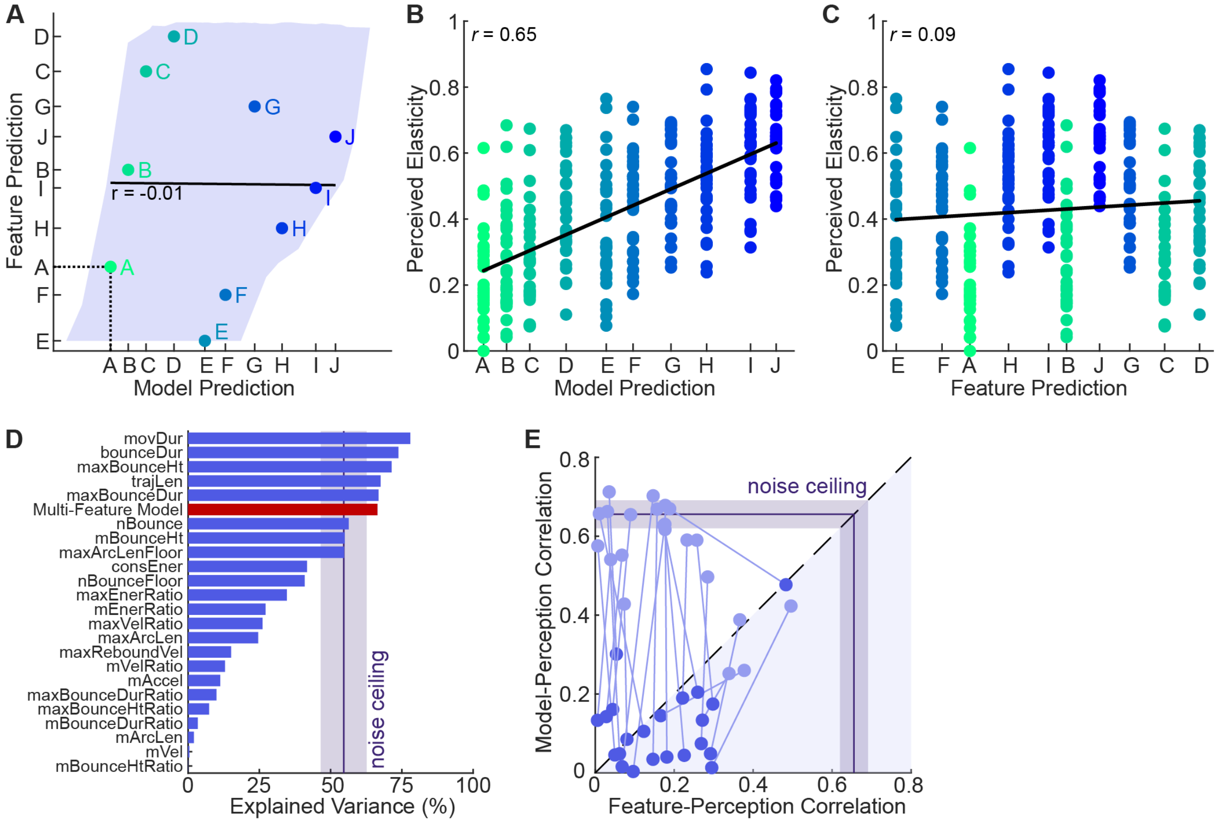


**Figure S6. Principle and results of the decorrelation experiment (Exp. 2), related to Figure 4. A)** Prediction of an example feature (consEner) plotted against the prediction of the multi-feature model. The blue-shaded area shows the convex hull around all possible stimuli (i.e., all 100,000 simulations of medium elasticity). Feature and multi-feature model were strongly correlated in this data set (*r* = 0.62). The colored dots show the ten stimuli that were selected for the experiment together with a regression line. Importantly, in this stimulus set, feature and multi-feature model are not correlated (*r* = -0.01). **B)** Perceived elasticity for the ten selected stimuli from 30 observers is plotted as a function of the multi-feature model prediction. The black line shows the linear fit. Model and perception correlate strongly (*r* = 0.65). **C)** This plot shows the same data as (B) but the position on the x-axis is ordered according to the feature prediction together with a linear fit. The correlation between feature and perception is significantly lower (*r* = 0.09). Thus, the multi-feature model is a better predictor of perceived elasticity than the consEner feature. **d)** The bar graph shows the explained variance of each predictor in terms of averaged perceived elasticity across *all* stimuli (i.e. not just for the separate sets of stimuli designed to decorrelate each feature from the model). This analysis suggests that movement duration is the best predictor of perceived elasticity. The noise ceiling shows the average (± 95%-CI) correlation of every participant with the average observer. **E)** This plot shows the correlation between perceived elasticity and the prediction of the multi-feature model against the correlation between perception and the feature prediction for 22 features (all except movDur; light blue dots). Additionally, the plot shows the partial model-perception and feature-perception correlations when controlling for the effect of movement duration. For most features, the high correlations between model and perception decrease drastically when controlling for the effect of movement duration. This suggests that high perception-model correlations are mainly an artifact of the high correlation between movement duration and feature model. The noise ceiling was calculated by correlating the pooled data with the average rating for each set of stimuli, i.e., for each feature. The purple line shows the average (± 95%-CI) across features.

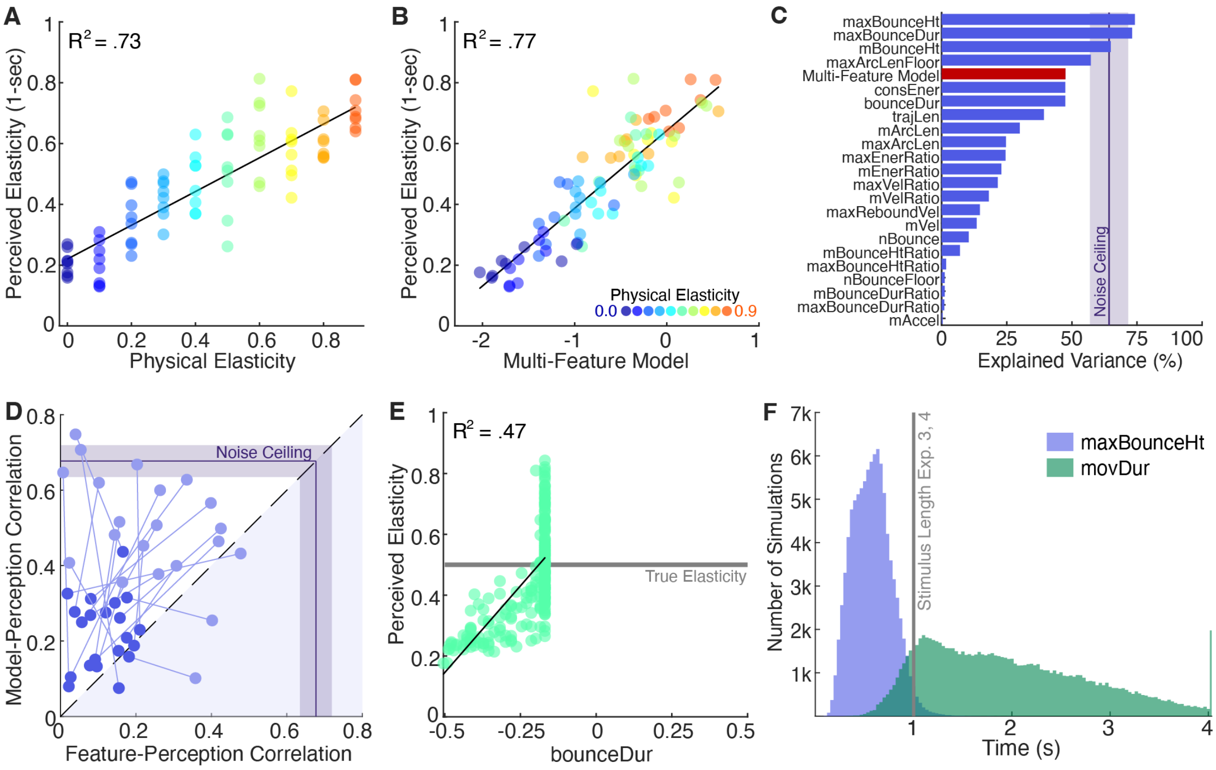


**Figure S7. Results of Experiments 3 and 4, related to Figure 5. A)** Average elasticity ratings for 1-sec movies as a function of physical elasticity together with a linear fit. Dots of the same color show simulations of the same elasticity but varying initial parameters. **B)** Average elasticity ratings for 1-sec movies as a function of the multi-feature model, together with a linear fit. Physical elasticity is indicated by the color of the dots. **C)** Bar graph shows the explained variance of each predictor in terms of averaged perceived elasticity across *all* stimuli in Experiment 4 (not just for the separate sets of stimuli designed to decorrelate a given feature from the model). This post hoc analysis suggests that the maximum bounce height (presumably as a proxy for the movement duration in full movies) is the best predictor of perceived elasticity in truncated movies. The noise ceiling shows the average (± 95%-CI) correlation of every participant with the average observer. **D)** This plot shows the correlation between perceived elasticity and the prediction of the multi-feature model against the correlation between perception and the feature prediction for 21 features (all except movDur and maxBounceHt; light blue dots). Additionally, the plot shows the partial model-perception and feature-perception correlations when controlling for the effect of maximum bounce height. For most features, the high correlations between model and perception decrease drastically when controlling for the effect of maximum bounce height. This suggests that high perception-model correlations are mainly an artifact of the high correlation between maximum bounce height and feature model. The noise ceiling was calculated by correlating the pooled data with the average rating for each set of stimuli, i.e., for each feature. The line shows the average (± 95%-CI) across features. **E)** Average elasticity ratings of Experiment 4 as a function of the bounce duration together with a linear fit. Elasticity ratings increase with an increase in bounce duration. However, because the movies are truncated bounce duration is only a diagnostic feature if it is notably shorter than one second. In this case, observers seem to rely on different cues. All stimuli had the same physical elasticity of 0.5. Thus, a physics-based model would predict all stimuli to appear equally elastic (grey line). **F)** Histograms of when the maxBounceHt and movDur are reached. 94.10% of all simulations reach their maxBounceHt within the first second.

**Table S1. Motion features with % variance explained in physical elasticity.**
Features< 5% were excluded from further analysis.

| % | Feature (acronym; unit) |
|---|---|
| 82.29 | Movement duration until the cube stopped moving. (movDur; sec) |
| 78.93 | Number of bounces from the floor, the ceiling and the walls. (nBounce) |
| 78.92 | Duration until the cube landed after the last bounce from any wall. (bounceDur; sec) |
| 77.87 | Number of bounces from the floor. (nBounceFloor) |
| 67.27 | Cumulative length of the motion trajectory. (trajLen; m) |
| 52.76 | Mean ratio of energy before and after a bounce. (mEnerRatio) |
| 51.91 | Mean acceleration over time. (mAccel; $m/s^2$) |
| 50.80 | Conserved energy over time. (consEner) |
| 45.85 | Maximum ratio of energy before and after a bounce. (maxEnerRatio) |
| 44.86 | Maximum length of bounce arcs from floor (maxArcLenFloor; m) |
| 41.80 | Mean ratio of incident to rebound velocity of all bounces. (mVelRatio) |
| 39.42 | Maximal ratio of incident to rebound velocity of all bounces. (maxVelRatio) |
| 36.51 | Maximal ratio of durations of consecutive bounces from the floor. (maxBounceDurRatio) |
| 35.46 | Maximal duration of individual bounces from the floor. (maxBounceDur; sec) |
| 35.21 | Maximal rebound velocity of bounces from every wall. (maxReboundVel; m/s) |
| 35.13 | Maximal ratio of bounce heights of two consecutive bounces from the floor. (maxBounceHtRatio) |
| 35.10 | Maximal height of bounces from the floor. (maxBounceHt; m) |
| 30.84 | Mean ratio of bounce durations of consecutive bounces from the floor. (mBounceDurRatio) |
| 23.42 | Mean ratio of bounce heights of two consecutive bounces from the floor. (mBounceHtRatio) |
| 16.25 | Maximal length of bounce arcs, i.e., trajectory between consecutive bounces. (maxArcLen; m) |
| 6.24 | Mean height of bounces from the floor. (mBounceHt; m) |
| 5.49 | Mean velocity over time. (mVel; m/s) |
| 5.25 | Mean length of bounce arcs, i.e., trajectory, between consecutive bounces. (mArcLen; m) |
| 4.06 | Mean length of bounce arcs from floor. (mArcLenFloor; m) |
| 1.86 | Difference between movement and bounce duration. (otherMotionDur; sec) |
| 0.77 | Mean duration of individual bounces from the floor. (mBounceDur; sec) |
| 0.15 | Mean height of the object over time. (mHeight; m) |
| 0.01 | Mean rebound velocity of bounces from all walls. (mReboundVel; m/s) |

**Table S2. List of motion features, their definitions and rationale, related to Table 1.**

| Feature (unit) | Acronym | Definition (Figure Number) | Rationale |
|---|---|---|---|
| **Features characterizing bounce events.** | | | |
| **Mean bounce height (m)** | mBounceHt | Bounce height was defined as the maximal height of the cube between two bounces from the floor (irrespective of whether another wall or the ceiling was hit in between).[1,2] We calculated mean and maximum across bounces. (S20) | Objects that are more elastic tend to bounce higher.[4,5] Humans might attend to the average or only to the highest bounce. |
| **Maximum bounce height (m)** | maxBounceHt | | |
| **Mean ratio of bounce heights** | mBounceHtRatio | Bounce height ratio was calculated as ratio of bounce heights of two consecutive bounces from the floor (unless the cube hit the ceiling in between as this would bias the results).[1,2] We calculated mean and maximum across bounces. (S21) | Heights of consecutive bounces tend to be more similar for cubes that are more elastic.[4,5] Humans might attend to the average or only to the maximal ratio. |
| **Maximum ratio of bounce heights** | maxBounceHtRatio | | |
| **Mean rebound velocity (m/s)*** | mReboundVel | Rebound velocity was defined as the velocity of the cube right after a bounce from any wall. [1,2] We calculated mean and maximum across bounces. (S26) | Objects that are more elastic tend to bounce off with higher speed.[4] Humans might attend to the average or only to the maximal rebound velocity. |
| **Maximum rebound velocity (m/s)** | maxReboundVel | | |
| **Mean of incident/ rebound velocity ratio** | mVelRatio | Incident velocity was defined as the velocity of the cube right before a bounce from any wall. The velocity ratio was defined as the ratio between incident to rebound velocity. [1,2] We calculated mean and maximum across bounces. (S27) | Objects that are more elastic tend to bounce off with larger speed relative to incident speed.[4,5] Humans might attend to the mean or only to the maximum velocity ratio. |
| **Maximum of incident/ rebound velocity ratio** | maxVelRatio | | |
| **Mean duration of individual bounces (sec)*** | mBounceDur | The duration of an individual bounce was defined as the duration between two bounces from the floor (irrespective of whether another wall or the ceiling was hit in between).[1,2] We calculated mean and maximum across bounces. (S22) | Bounces of elastic objects tend to take longer.[4,5] Humans might attend to the mean or only to the maximum duration of individual bounces. |
| **Maximum duration of individual bounces (sec)** | maxBounceDur | | |

| | | | |
|---|---|---|---|
| **Mean ratio of bounce durations** | mBounceDurRatio | Bounce duration ratio was calculated as ratio of bounce durations of two consecutive bounces from the floor (unless the cube hit the ceiling in between as this would bias the results).[1,2] We calculated mean and maximum across bounces. (S23) | Durations of consecutive bounces are more similar for cubes that are more elastic.[4] Humans might attend to the mean or only to the maximum ratio of durations of individual bounces. |
| **Maximum ratio of bounce durations** | maxBounceDurRatio | | |
| **Mean length of bounce arcs (m)** | mArcLen | The length of a bounce arc was defined as the cumulative length of the trajectory between two consecutive bounces from any wall.[1,2] We calculated mean and maximum across bounces. (S24) | Objects that are more elastic tend to travel a longer distance per bounce. Humans might attend to the mean or only to the maximum length of bounce arcs. |
| **Maximum length of bounce arcs (m)** | maxArcLen | | |
| **Mean length of bounce arcs from floor (m)*** | mArcLenFloor | The length of a bounce arc was defined as the cumulative length of the trajectory between two consecutive bounces from the floor (irrespective of whether another wall or the ceiling was hit in between).[1,2] We calculated mean and maximum across bounces. (S25) | Objects that are more elastic tend to travel a longer distance per bounce. Humans might attend to the mean or only to the maximum length of bounce arcs. |
| **Maximum length of bounce arcs from floor (m)*** | maxArcLenFloor | | |
| **Mean energy ratio per bounce** | mEnerRatio | Energy ratio was defined as the ratio of energy[3] before and after a bounce from any wall.[1,2] We calculated mean and maximum across bounces. (S19) | The more elastic an object, the less kinetic and potential energy is transformed into other forms during a bounce (thus the object bounces off higher and faster). Humans might attend to the mean or only to the maximum of the energy ratio per bounce. |
| **Maximum energy ratio per bounce** | maxEnerRatio | | |
| | **Features integrating statistics of the motion over time.** | | |
| **Mean velocity (m/s)** | mVel | Euclidian distance travelled per second averaged over time.[1] (S14) | Objects that are more elastic tend to move with higher average speed. |
| **Mean acceleration ($m/s^2$)** | mAccel | Derivative of velocity over time averaged over time.[1] (S15) | Objects that are more elastic tend to decelerate more slowly. |
| **Mean height (m)*** | mHeight | Height of the cube averaged over time.[1] (S16) | Objects that are more elastic tend to spend more time up in the air. |
| **Trajectory length (m)** | trajLen | Cumulative sum of Euclidian distance travelled per frame.[1] (S17) | Objects that are more elastic tend to travel longer distances. |
| **Movement duration (sec)** | movDur | Duration until the cube stopped moving, i.e. its velocity was < 0.003 m/s. (S10) | Objects that are more elastic tend to move for a longer duration. |

| | | | |
|---|---|---|---|
| **Duration until the last bounce (sec)** | bounceDur | Duration until the cube lands after the last bounce from any wall.[1,2] (S11) | Objects that are more elastic tend to bounce for a longer duration. |
| **Duration of other (non-bounce) motion (sec)*** | otherMotionDur | Duration during which the cube is moving, but not bouncing (e.g., rolling, sliding, tumbling) defined as the difference between motion and bounce duration. (S11) | Objects that are more elastic tend to move for a longer duration even after the last bounce, e.g. rolling over the floor (without lifting up). |
| **Number of all bounces** | nBounce | Number of bounces from the floor, the ceiling and the walls.[1,2] (S12) | Objects that are more elastic tend to bounce more often from the floor, the ceiling and the walls. |
| **Number of bounces from the floor** | nBounceFloor | Number of bounces from the floor.[1,2] (S13) | Objects that are more elastic tend to bounce more often from the floor. |
| **Conserved energy over time** | consEner | Conserved energy was defined as the average energy[3] during the last flight phase given the average energy during the first flight phase ($e_{end}/e_{first}$) normalized by the time passed.[1,2] (S18) | The conserved energy over a given duration is higher for objects that are more elastic. |

*This feature was not included in the final model because its explained variance < 5% in terms of physical elasticity.

[1] Only for frames in which cube was in motion, i.e. velocity above threshold of 0.003 m/s.

[2] A bounce was detected if one of the cube's corners or its CoM (± ½ edge length) was in contact with a wall, preceded and followed by all points of the cube being in the air. The position of the CoM and the eight corners of the cube were interpolated between frames to detect bounces outside the frame rate.

[3] Energy was approximated as the potential maximal speed of fall, which is proportional to the sum of kinetic and potential energy. We calculated the energy *e* based on the velocity *v* and height *h* of the cube (i.e., ignoring its rotation and deformation): $e = v + \sqrt{2gh}$, where *g* denotes gravity (9.81 m/s$^2$).

[4] Proposed by Warren et al[28]

[5] Proposed by Nusseck et al[29]
